# Inhibition of DNA2 nuclease as a therapeutic strategy targeting replication stress in cancer cells

**DOI:** 10.1101/107383

**Authors:** Sandeep Kumar, Xiangdong Peng, James M. Daley, Lin Yang, Jianfeng Shen, Nghi Nguyen, Goeun Bae, Hengyao Niu, Yang Peng, Hui-Ju Hsieh, Lulu Wang, Chinthalapally Rao, Clifford C. Stephan, Patrick Sung, Grzegorz Ira, Guang Peng

## Abstract

Replication stress is a characteristic feature of cancer cells, which is resulted from sustained proliferative signaling induced by activation of oncogenes or loss of tumor suppressors. In cancer cells, oncogene-induced replication stress manifests as replication-associated lesions, predominantly double-strand DNA breaks (DSBs). An essential mechanism utilized by cells to repair replication-associated DSBs is homologous recombination (HR). In order to overcome replication stress and survive, cancer cells often require enhanced HR repair capacity. Therefore, the key link between HR repair and cellular tolerance to replication-associated DSBs provides us with a mechanistic rationale for exploiting synthetic lethality between HR repair inhibition and replication stress. Our studies showed that DNA2 nuclease is an evolutionarily conserved essential component of HR repair machinery. Here we demonstrate that DNA2 is indeed overexpressed in pancreatic cancers, one of the deadliest and more aggressive forms of human cancers, where mutations in the KRAS are present in 90%-95% of cases. In addition, depletion of DNA2 significantly reduces pancreatic cancer cell survival and xenograft tumor growth, suggesting the therapeutic potential of DNA2 inhibition. Finally, we develop a robust high-throughput biochemistry assay to screen for inhibitors of the DNA2 nuclease activity. The top inhibitors were shown to be efficacious against both yeast Dna2 and human DNA2. Treatment of cancer cells with DNA2 inhibitors recapitulates phenotypes observed upon DNA2 depletion, including decreased DNA end resection and attenuation of HR repair. Similar to genetic ablation of DNA2, chemical inhibition of DNA2 selectively attenuates the growth of various cancer cells with oncogene-induced replication stress. Taken together, our findings open a new avenue to develop a new class of anti-cancer drugs by targeting druggable nuclease DNA2. We 4, 16. In propose DNA2 inhibition as new strategy in cancer therapy by targeting replication stress, a molecular property of cancer cells that is acquired as a result of oncogene activation instead of targeting undruggable oncoprotein itself such as KRAS.

## Introduction

The finding that poly(ADP-ribose) polymerase (PARP) inhibitors, a new class of DNA repair inhibitors, specifically kill cancer cells with BRCA1 or BRCA2 mutations but are less cytotoxic to normal cells highlighted the promise of DNA repair inhibitors for targeted cancer treatment ^7, 21^. This finding also provides proof of the principle that synthetic lethality interactions in the DNA repair network can be exploited for targeted cancer therapy. Early in the process of tumorigenesis, genetic alterations such as activation of oncogenes and loss of tumor suppressor genes are implicated in inducing replication stress by providing premalignant cells with excessive growth signals ^4, 16^. In cancer cells, oncogene-induced replication stress manifests as a high level of DNA double-strand breaks (DSBs) due to stalling and collapse of DNA replication forks ^22, 37^. A dramatic increase of replication stress and spontaneous DNA damage in cancer cells, which renders them more dependent on DNA repair for survival ^18^. Among all DNA repair pathways, homologous recombination-mediate DNA repair (HR) is an essential mechanism utilized by cancer cells to repair replication-associated DSBs and thereby to overcome replication stress and survive. The key link between HR repair and cellular tolerance to replication-associated DSBs provides us with a mechanistic rationale for exploiting synthetic lethality between HR repair inhibition and replication stress in cancer cells.

Moreover, owing to their hyper-proliferative state, cancer cells are more vulnerable to killing by DNA damaging agents. Common cancer treatment strategies use untargeted radiomimetic chemotherapy or radiotherapy that induce a broad range of DNA lesions including DSBs. The ability of cancer cells to repair DNA damage lowers therapeutic efficacy, thus the simultaneous use of a DNA repair inhibitor holds great promise in sensitizing cancer cells to conventional chemo/radiotherapy. Inhibitors that target enzymes mediating base excision repair, nucleotide excision repair and other DNA repair pathways have been developed and are being evaluated in clinical trials ^11, 17, 32^. Therefore, identification of new HR repair inhibitors can open new avenues to cancer therapies ^2, 3, 9, 16, 23, 30^.

An evolutionarily conserved protein DNA2 possesses 5’ flap endonuclease and 3’-5’ helicase activities and it plays an important role in DNA damage repair, homologous recombination, and DNA replication. The nuclease activity of DNA2 has several well-documented cellular functions, while the biological role of its helicase activity remains enigmatic. Specifically, DNA2 mediates the resection of the 5’ strand at DNA double strand break ends ^12, 25, 26, 28, 34, 39, 40, 51^, an early step in homologous recombination, and also at stalled and regressed replication forks ^29, 46^. Dna2 in fungi processes long 5’ DNA flaps during the maturation of Okazaki fragments in lagging strand DNA synthesis ^1, 8^, while human DNA2 plays an essential but as yet undefined role in DNA replication ^20^. Human DNA2 has an additional function in mitochondrial DNA maintenance ^50^. Finally, the unstructured and regulatory N-terminal domain of yeast Dna2 was recently shown to regulate DNA damage response by activating the Mec1 kinase ((Mec1 is orthologus to human DNA damage checkpoint kinase ATR) ^33^. Among these important biological functions of DNA2, the nucleolytic processing of the 5’ strands at spontaneous DSBs or regressed replication forks seems to be the most relevant in alleviating replication stress. Consistent with this notion, DNA2 is overexpressed via gene duplication in yeast upon replication stress caused by the *mec1-21* mutation ^48^. Interestingly, similar to this observation in yeast cells, studies from our group and others have demonstrated that DNA2 is overexpressed in a broad spectrum of malignancies including breast cancer, ovarian cancer, lung cancer, prostate cancer, glioblastoma, pancreatic cancer, leukemia, colon cancer, melanoma, Lynch Syndrome, and hereditary nonpolyposis colorectal cancer ^19, 42, 45^. DNA2 overexpression is needed for the tolerance of increased replication stress induced by oncogene activation such as H-Ras or cyclin E in osteosarcoma and breast cancer cells, and partial depletion of DNA2 reduces the tumorigenicity of breast cancer cells ^42^. These results have raised the prospect of targeting DNA2 in cancer therapy.

Since the increased nuclease activity of DNA2 is required in yeast and cancer cells to overcome replication stress and DNA damage, for the first time, we developed a robust biochemistry-based screening method for chemical inhibitors of DNA2. We conducted initial screen with nearly 50,000 compounds and we tested efficacy of several candidate inhibitors in pancreatic and breast cancer cell lines with oncogene-induced replication stress. Treatment of cancer cells with the top DNA2 inhibitor recapitulates many phenotypes observed upon DNA2 depletion, and attenuates the growth of various cancer cells. Taken together, in this study, we developed a feasible high-throughput chemical screen assay to identify novel DNA repair inhibitors and more importantly we provided a new synthetic lethal strategy based on inhibition of HR repair by DNA2 inhibitors in cancer cells with oncogene-induced replication stress.

## Results

### Sensitivity of pancreatic cancer cells to DNA2 depletion

Pancreatic cancer is particularly deadly due to the lack of an effective treatment regimen. Here, we investigated the expression levels and therapeutic potential of DNA2 depletion in pancreatic cancers, where 90-95% cases are associated with an activating mutation in the K-Ras oncogene ^13, 24, 38^. Analysis of gene expression patterns in pancreatic cancers in the Oncomine database shows that DNA2 is significantly upregulated ^42^. Consistently, an elevated DNA2 protein level was observed in pancreatic cancer patient specimens (Figure 1a). Moreover, patients with chronic pancreatitis expressed DNA2 at an elevated level in their pancreatic tissue (Figure 1a). To establish the relationship between the K-Ras oncogene and DNA2 expression, we analyzed a transgenic *K-Ras^G12D^* pancreatic cancer mouse model. Similar to pancreatic cancer patient specimens, the DNA2 protein level was greatly increased in *K-Ras^G12D^*-activated early pancreatic cancer lesions, suggesting that DNA2 overexpression is an early event during tumorigenesis (Figure 1b). Importantly, the depletion of DNA2 impaired the growth and survival of cells from two pancreatic cancer lines, and also led to increased senescence and apoptosis of these cells (Figure 1c, Supplementary Figure S1a). Consistent with in vitro cell culture findings, DNA2 knockdown significantly reduced tumor growth in the xenograft model (Supplementary Figure S1b). Furthermore, our analyses showed increased staining of p-CHK1, a key molecule in DNA damage signaling, and reduced staining of Ki67, a maker of cell proliferation, in DNA2-depleted cells likely due to endogenous DNA damage (Supplementary Figure S1c). Thus, DNA2 overexpression is a common theme in many cancer types including pancreatic cancer and inhibition of DNA2 reduces pancreatic cancer cell survival both *in vitro* and *in vivo*.

**Figure 1.**
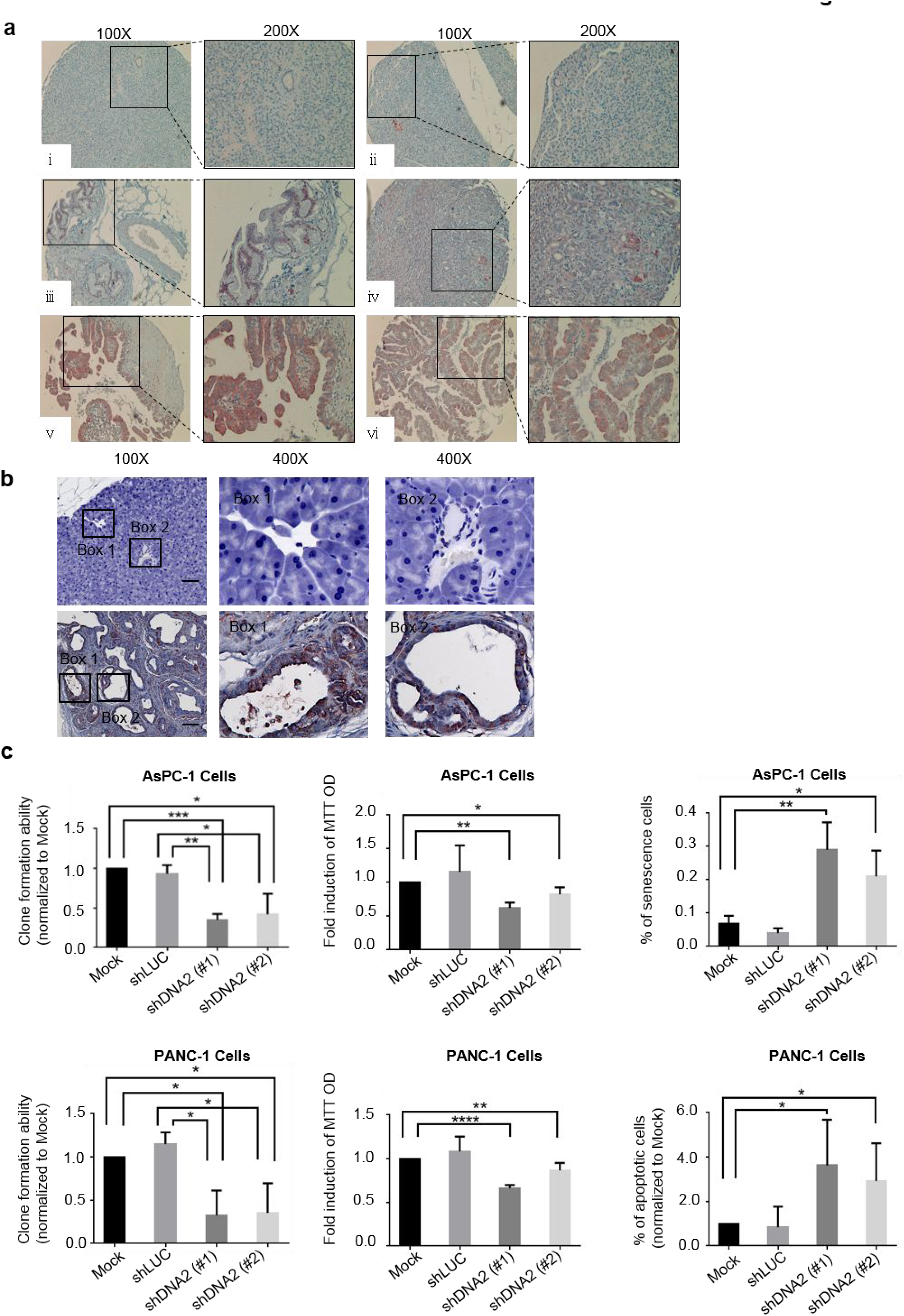
DNA2 is overexpressed in pancreatic cancer lesions. (**a**) Immunohistochemistry (IHC) staining of DNA2 in representative human pancreatic lesion specimens. Two cases are presented and the lesions (boxed) are also shown at a higher magnification. (ⅰ, ⅱ): normal pancreatic tissue; (ⅲ, ⅳ): hyperplasia in pancreatic tissue with chronic pancreatitis; (ⅴ, ⅵ) pancreatic carcinoma. (**b**) IHC staining of DNA2 in wild type mouse pancreatic tissue (top) and in LSL-K-Ras^G12D^ mouse pancreatic cancer tissue (bottom). Scale bar, 100 μm. Segments (boxed) of the images are also shown at a high magnification. (**c**) DNA2 depletion affects the survival of pancreatic cancer cells. Pancreatic cancer cell lines AsPC-1 and PANC-1 were stably transfected with shRNA vectors against two non-overlapping sequences in DNA2 or scramble control (Mock) (see also Supplementary Figure S1). (Left) Clonogenic assay, cell survival scored 10-15 days after seeding. (Middle) MTT assay, cells proliferation measured 72 hrs after seeding. (Right) Senescence assay with β-galactosidase staining and apoptosis assay using annexin V staining performed 72 hours after DNA2 knockdown. Each value represents the mean ± SEM from three independent experiments (* p<0.05; ** p<0.01; *** p<0.005; **** p<0.001).

### Isolation of yeast Dna2 inhibitors

We have presented evidence that DNA2 overexpression alleviates the impaired proliferation of U2OS and MCF10A cells after an activation of either H-Ras or cyclin E ^42^, suggesting that DNA2 could be an effective target in cancer therapy. This prompted us to develop a simple screen for chemical inhibitors of DNA2. The screen entailed the use of oligonucleotide (dT)_30_ having a reporter fluorescent moiety, 6-FAM^TM^ on the 5’ end, and the dark quenching group, Iowa Black^®^ FQ, on the 3’ end. Additionally a second dark quencher ZEN^TM^ was placed between the 9th and 10th base (Figure 2a). The fluorescence emitted by 6-FAM^TM^ is quenched by FRET, but it increases when DNA is degraded by Dna2. In the initial screen, compounds were tested at a fixed concentration of inhibitors (33.33 µM) with a sub-saturating amount (0.09 nM) of Dna2. The inhibitory activity of the compounds was determined as a reduction in fluorescent signal compared to the DMSO only control. The primary screen hits were selected based on a ≥50% inhibition of the Dna2 activity. We screened ~50,000 chemical compounds from six different libraries (Supplementary Table S1) to identify chemical inhibitors of the Dna2 nuclease activity. Since we could not purify human DNA2 in amounts sufficient for a high throughput screen, the initial screen was performed with yeast Dna2 (yDna2), followed by testing top candidate compounds with human DNA2 (hDNA2). The DNA2 nuclease domain is well conserved among eukaryotes, and therefore we reasoned that inhibitors of yDna2 will likely also work with the human counterpart. DNA2 nuclease efficiently cleaves the single stranded DNA substrate used here (Figure 2b and ^44^), and inhibitors are expected to block substrate cleavage. First, we determined the concentration of yeast and human DNA2 and of the control T5 nuclease needed to generate maximum fluorescence (Figure 2c), and we optimized the screen reaction (Z’ value of 0.8) in a robotic system with a 384-well plate setup to provide high throughput. The first screening round identified 184 compounds, corresponding to a hit rate of ~0.4%. These compounds were subject to re-screening to confirm their efficacy and also to eliminate those compounds that also inhibit T5 nuclease. We were able to narrow yDna2-specific candidate compounds down to 39 (Supplementary Table S2).

**Figure 2.**
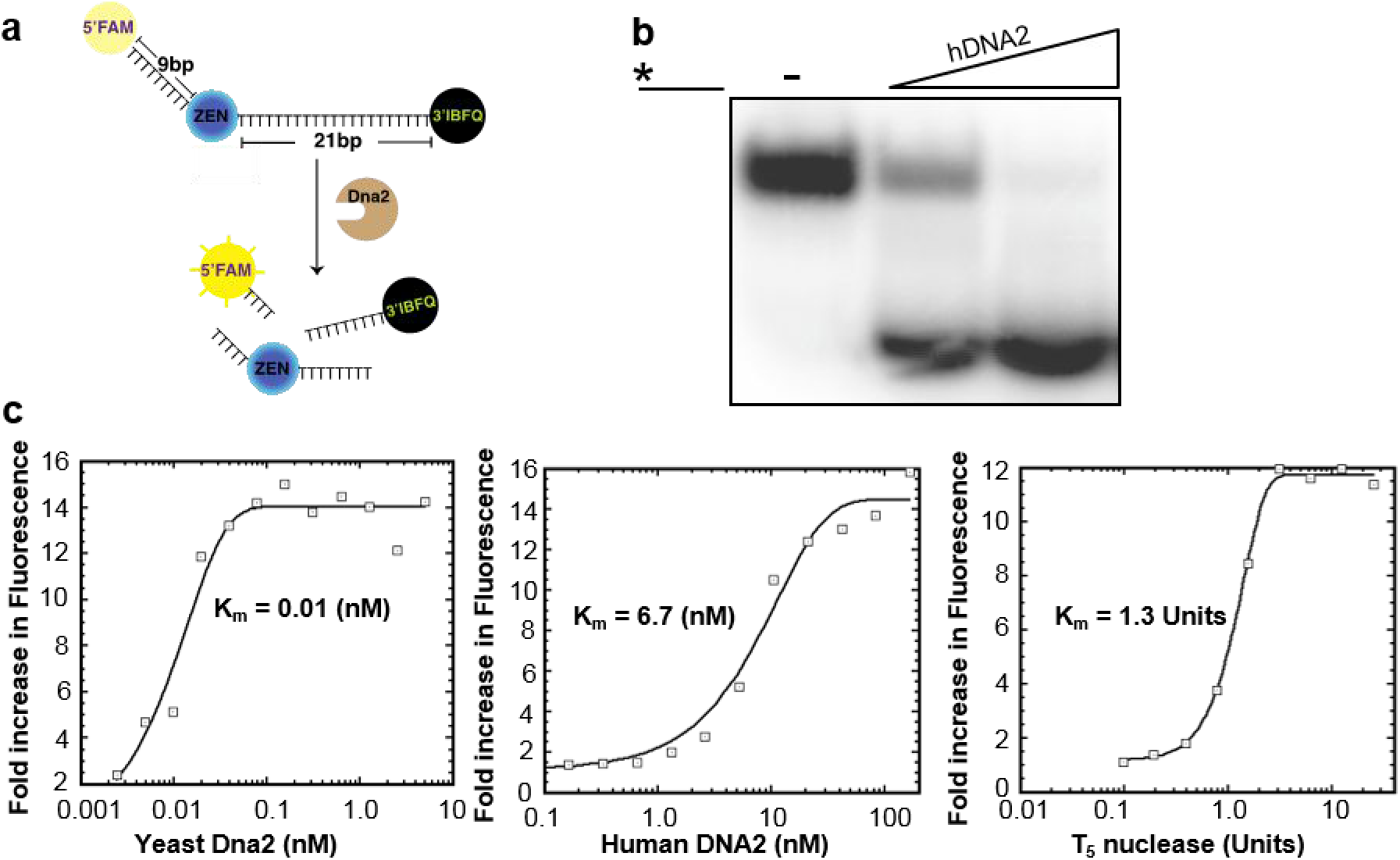
Assay optimization for high throughput screen for DNA2 inhibitors. (**a**) Graphical representation of the screen assay. The 30 nucleotides long ssDNA substrate is labeled with three dyes: a fluorophore (6-FAM^TM^) at the 5’end, and two dark quenchers, internal ZEN^TM^ (between 9th and 10th base) and Iowa Black^®^ FQ at the 3’ end. The extent of the reaction was determined by measuring increase of fluorescence at wavelength of 520 nm. (**b**) hDNA2 (0, 0.5, or 1 nM) was incubated with radiolabeled ssDNA (2.5 nM). The radiolabel is denoted by *. (**c**) Degradation of triply labeled ssDNA (100 nM) by yDna2 (0.002 to 5.0 nM), hDNA2 (0.00016 to 166 nM) and T5 nuclease (0.01 to 10 Units).

We note that a number of the compounds listed in Supplementary Table S3 inhibited the activity of both yDna2 and T5 and are therefore non-specific. Interestingly, many of these nonspecific inhibitors interact with DNA and are already used in cancer therapy, including multiple drugs known to inhibit type II topoisomerases (e.g. Daunorubicin, Doxorubicin, Mitoxantrone). Here, we have provided evidence that these compounds also affect the activity of Dna2. Thus, in addition to inducing DSBs by blocking topoisomerase II, it is likely that these drugs affect the processing and repair of these DSBs by nucleases.

### Cellular screen for the most potent DNA2 inhibitors

The biological effectiveness of the 39 compounds selected by re-screening was determined by measuring their half maximal inhibitory concentration (IC_50_) biochemically and in a cell-based system using the 384-well platform (Supplementary Table S2). We used the cancer cell line U2OS, as cells of this lineage overexpress DNA2 significantly and are sensitive to partial DNA2 depletion ^42^. We conducted the viability assay in duplicate on three separate occasions using a concentration range of 10 pM to 10 µM for each compound. Cell proliferation was determined using DAPI staining and microscopy. About ten compounds showed IC_50_ values between <1 μM and 10 μM both in cells and in vitro (Supplementary Table S2), and two of them, NSC-5195242 and NSC-105808 (Figures 3a and b) were selected for further analysis.

**Figure 3.**
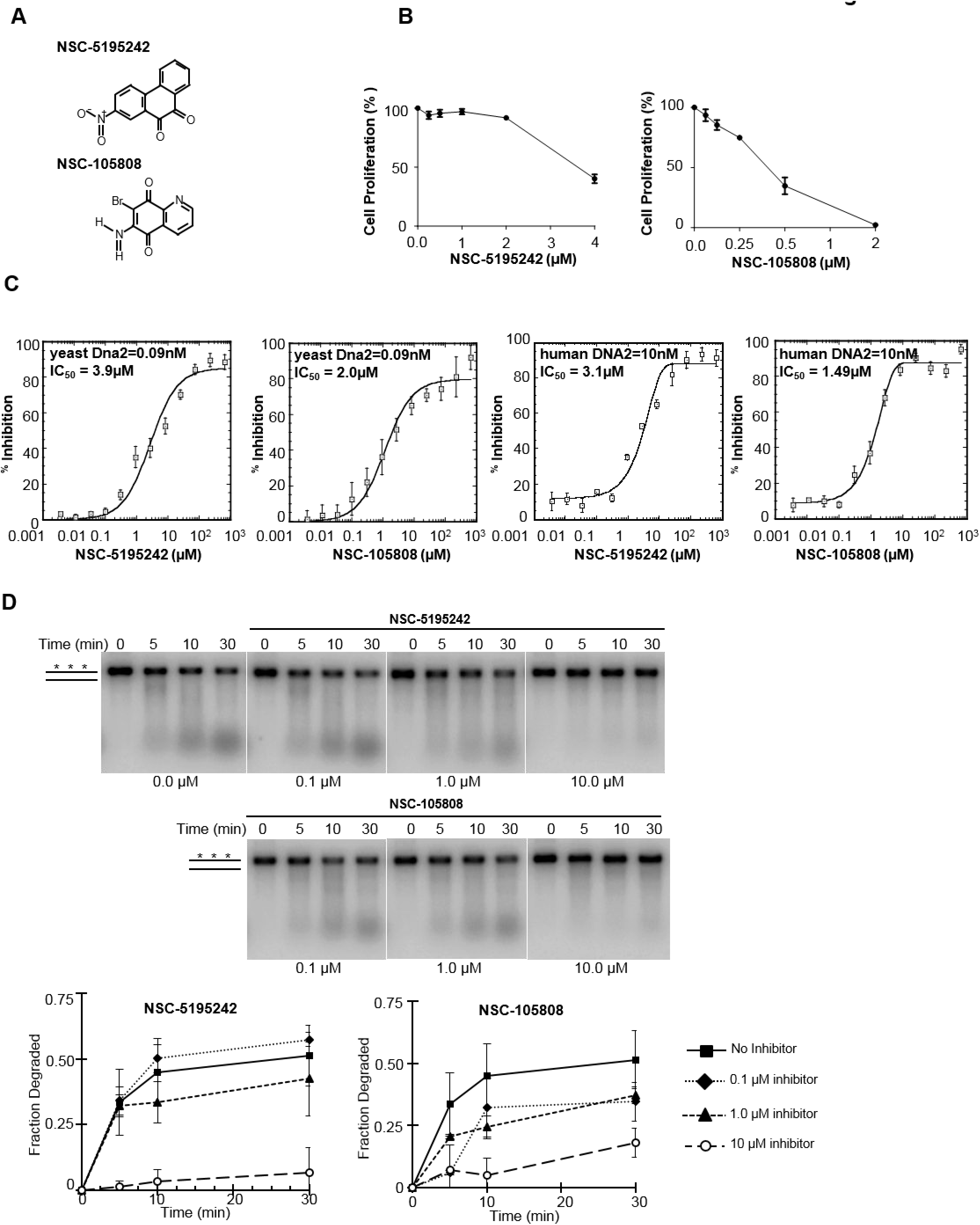
Validation of the top two inhibitors identified from the high throughput screen. (**a**) Chemical structure of NSC-5195242 (2-nitrophenanthrene-9,10-dione) and NSC-105808 (6-amino-7-bromoquinoline-5,8-dione). (**b and c**) Dose response curves for indicated compounds, cell based (U2OS cell line) (**b**) and in vitro (**c**). The IC_50_ value and the protein concentration used are indicated. Triple labeled substrate (0.1 µM) as in Figure 2a was used. The concentration range of inhibitors was 0.003 to 666.67 µM. (**d**) Degradation of radiolabeled 2 kb dsDNA (0.5 nM ends) with hDNA2 (5 nM), BLM (5 nM), and RPA (100 nM) and the indicated concentrations of NSC-5195242 and NSC-105808. The asterisks denote the radiolabel. Plotted are the average data from three independent experiments and the error bars represent one standard deviation.

### Inhibitory effect of NSC-5195242 and NSC-105808 on human DNA2

The initial inhibitor screen was done with yDna2, thus it was important to test whether NSC-5195242 and NSC-105808 would affect the activity of hDNA2. hDNA2, purified as described ^14^, was first tested using the triply labeled substrate as above (Figure 2a). We found a similar inhibitory effect on hDNA2 and comparable IC_50_ values as determined for yDna2 (Figure 3c). Furthermore, we tested the impact of these two compounds in a reconstituted 5’ DNA end resection assay including hDNA2, BLM, and RPA as described ^14^ and in 5’ flap processing by hDNA2. Both compounds inhibited resection in the reconstituted system (Figure 3d) and also 5’ flap processing (Supplementary Figure S2). Considering that compound NSC-105808 shows a lower IC_50_ in vitro and biologically, further tests were focused on this compound. We found that NSC-105808 has no effect on ATPase activity of DNA2, nor the ability of BLM to unwind DNA, nor does it affect DNA binding by DNA2 (Supplementary Figure S2). Furthermore, NSC-105808 does not inhibit the nuclease activity of EXO1 (Supplementary Figure S2) which can also catalyze DNA ends resection. Together, these data provide further evidence that NSC-105808 specifically inhibits the nuclease activity of DNA2.

### NSC-105808 interferes with DNA end resection and homologous recombination in cells, and suppresses the cisplatin sensitivity of FANCD2-/- cells

To test whether NSC-105808 inhibits proliferation of U2OS cells by targeting DNA2, we first examined whether the ectopic overexpression of DNA2 could reduce the negative effect of NSC-105808 on cell growth. An approximately 1.5-2.0-fold increase in DNA2 protein level was able to attenuate the negative impact of NSC-105808 on cell proliferation, supporting the idea that DNA2 is the target of NSC-105808 in cells (Supplementary Figure S3). DNA2 functions in DSB end resection and homologous recombination (e.g.^25, 26, 42^), thus we asked whether NSC-105808 interferes with these biological processes. Similar to DNA2 knockdown, NSC-105808 (24 hr treatment) significantly reduced DSB repair by gene conversion or single stand annealing (SSA) as determined by using two GFP-based reporter assays (Figure 4a, Supplementary Figures S4-6 and ^42^). The treatment with NSC-105808 had only a mild impact on cell cycle distribution at the highest tested concentration (0.6 μM) with a slight decrease of G1 and increase of G2 cells, but with no impact on the cellular level of DNA2 protein (Supplementary Figures S7 and 8). We note that longer (>48 hr) exposure to NSC-105808 causes cell death (Figure 3b). To test DSB end resection, we followed phosphorylation of the ssDNA-binding protein RPA (p-RPA) as described previously ^43^. Cells were treated with the topoisomerase I inhibitor Camptothecin (CPT), which induces DNA nicks, reversed replication forks and single-ended DSBs. As shown in Figure 4b, NSC-105808 treatment significantly attenuated the accumulation of CPT induced p-RPA, suggesting an impairment of DSB end resection. We then confirmed this result by conducting chromatin fractionation and observed that NSC-105808 reduced p-RPA accumulation in the chromatin fraction with a concomitant enhancement in the γ-H2AX level, reflective of an accumulation of unrepaired DSBs (Figure 4c). Altogether, the above results provide evidence that NSC-105808 interferes with DSB resection and homologous recombination in cells.

**Figure 4.**
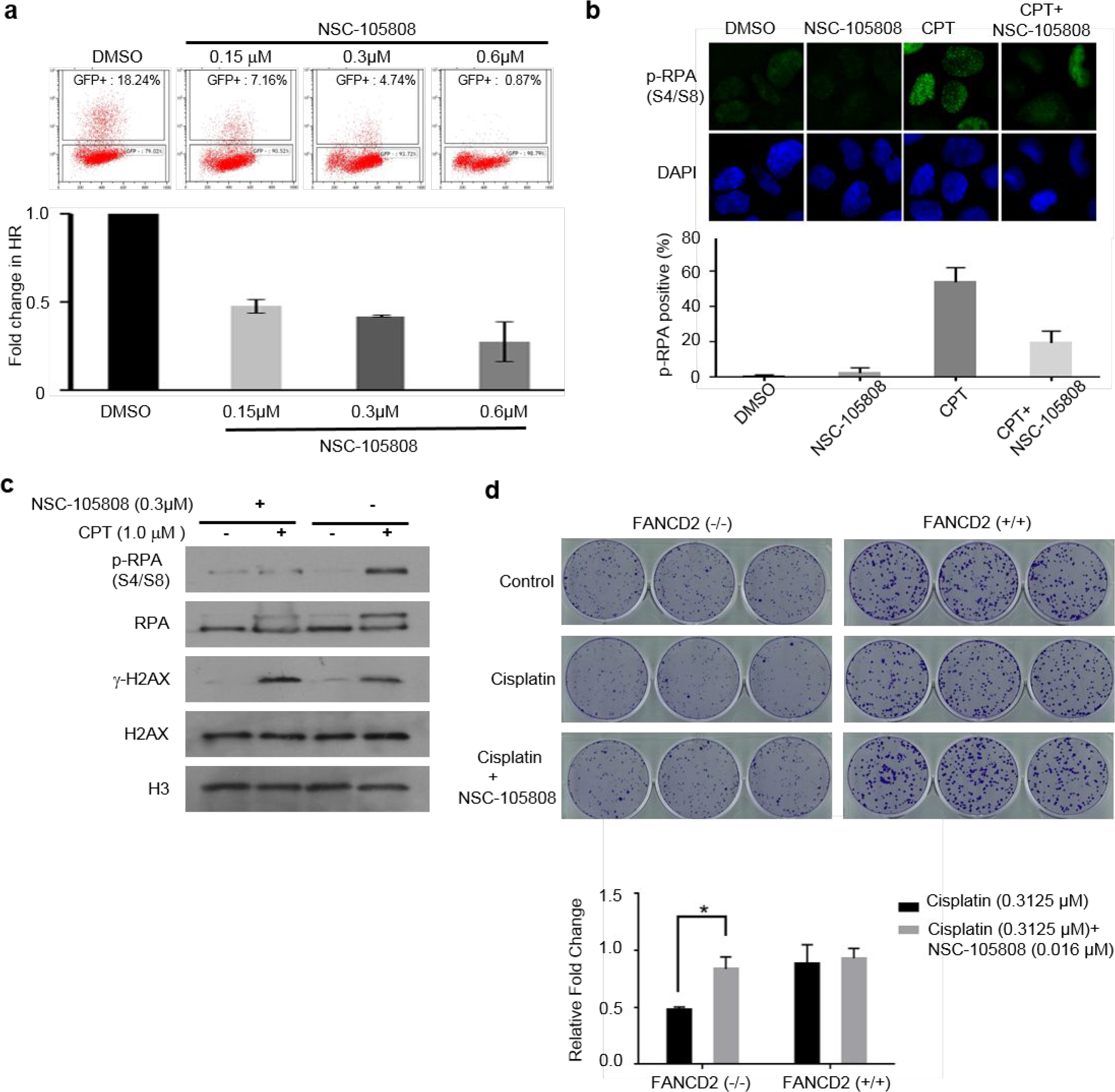
NSC-105808 inhibits HR repair and DSB end resection. (**a**) Analysis of HR efficiency with the DR-GFP assay. Representative images of flow cytometry analysis and bar graph showing percentage of GFP cells from at least three independent experiments. Each value represents the mean ± SD. (**b and c**) Analysis of RPA phosphorylation in U2OS cells treated with NSC-105808 (2 hr, 0.3 μM) and/or CPT (2 hr, 1 μM). (**b**) Representative images of cells subjected to immunofluorescent staining of p-RPA and bar graph showing percentage of cells with positive p-RPA staining from three independent experiments. Each value represents the mean ± SD. (**c**) Western blot analysisof chromatin fraction with the indicated antibodies. (**d**) Analysis of colony formation by FANCD2 (+/+) and FANCD2 (−/−) cells treated with cisplatin and/or NSC-105808. Each value is relative to the untreated control group of each cell line. The graph represents the mean ± SEM from three independent experiments. * p<0.01.

We wished to further verify the DNA2 specificity of NSC-105808 in cells. Previous work revealed that DNA2 depletion in FANCD2-deficient cells partially suppresses the sensitivity of these cells to cisplatin ^31^. As shown in Figure 4d, FANCD2-/- cells were more sensitive to cisplatin treatment compared with control cells, and, importantly, a very low concentration of NSC-105808 lessened the cisplatin sensitivity of these cells. Together, our results are consistent with the premise that DNA2 is the target of NSC-105808 in cells.

### Oncogene induction renders cells susceptible to NSC-105808

DNA2 overexpression, often observed in cancer cells, has been proposed to alleviate oncogene-induced replication stress. Increased sensitivity of cancer cells to DNA2 depletion ^42^ can likely be explained by an increased amount of substrates that are processed by DNA2 nuclease/helicase. We therefore asked whether NSC-105808 can sensitize cancer cells to oncogene-induced replication stress. Initially, we compared the impact of our DNA2 inhibitor in pancreatic or breast cancer cells versus control cells (Figures 5a and b). Transformed pancreatic cancer cell lines (AsPC-1, PANC-1) were more sensitive to NSC-105808 when compared to non-transformed pancreatic ductal epithelial cells (DT-PD59). Similarly, transformed breast cancer cells (Hs578T, BT549) were more sensitive than MCF10A non-transformed breast epithelial cells to the DNA2 inhibitor.

**Figure 5.**
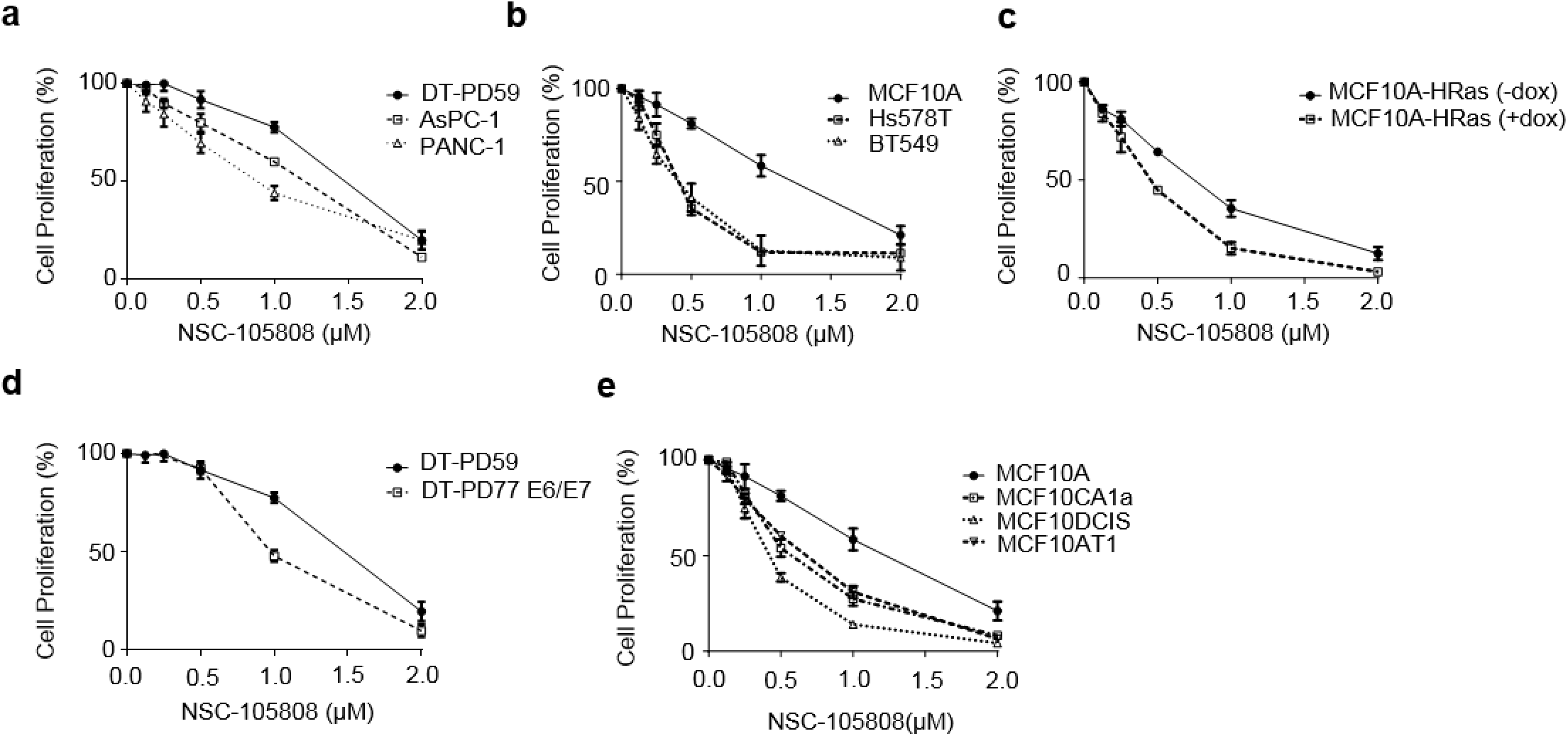
Oncogene induction enhances sensitivity of cancer cells to NSC-105808. (**a-e**). Graphs showing proliferation of indicated cell lines in response to a 48 hr treatment with the indicated concentrations of NSC-105808. Proliferation relative to cells treated with vehicle alone was determined by the MTT assay. Each value represents the mean ± SEM from three independent experiments. (**a**) Proliferation of control pancreatic epithelial cells DT-PD59 and transformed pancreatic cancer cells AsPC-1 and PANC-1. (**b**) Proliferation of normal breast epithelial cells MCF10A and breast cancer cells Hs578T and BT549. (**c**) Proliferation of MCF10A cells with or without H-Ras expression. H-Ras was induced with doxycycline (2 μg/ml). (**d**) Proliferation of control pancreatic DT-PD59 cells and transformed DT-PD77 cells with E6/E7 oncoprotein activation. (**e**) Proliferation of MCF10A breast cells without and with H-Ras activation (AT1, DCIS, CA1a).

Three separate cell models were then used to test whether NSC-105808 can target cancer cells with oncogene activation (Figures 5c-e). First, we generated an MCF10A cell line, an immortalized non-transformed breast epithelial cell line, with inducible H-Ras expression. Induction of H-Ras expression is shown in Supplementary Figure S9a. Upon activation of H-Ras, cells exhibited increased sensitivity to the DNA2 inhibitor (Figure 5c). Second, we used the E6/E7 oncoprotein to induce replication stress in non-transformed pancreatic ductal cells (DT-PD59) as previously described ^6^. Again, sensitivity to NSC-105808 was higher in cells with oncoprotein-induced replication stress (Figure 5d). In addition, we also established an inducible K-Ras expression cell line using non-transformed pancreatic ductal HPDE cells (Supplementary Figure S9a). Both MTT and colony formation assays showed that activation of K-Ras expression sensitized HPDE cells to NSC-105808 treatment (Supplementary Figure S9b and c). Lastly, we tested a series of epithelial cell lines that represent different stages of breast cancer progression ^15^: control non-tumorigenic MCF10A cells; premalignant/tumorigenic MCF10AT1 cells, derived from MCF10A cells by the expression of oncogenic H-Ras; tumorigenic/locally invasive MCF10-DCIS cells, and tumorigenic/metastatic MCF10CA1a cells, cloned from xenograft lesions induced by premalignant MCF10AT1 cells. Consistent with other observations, the three cell lines with oncogene activation were more sensitive to DNA2 inhibitor treatment compared to the MCF10A control (Figure 5e). Thus, in different cell models, oncogene activation sensitizes cells to NSC-105808.

## Discussion

During tumorigenesis, the activation of an oncogene or loss of a tumor suppressor gene leads to excessive growth signals and replication stress. Notably, about 30% of all cancers have an activating mutation in Ras and the vast majority (up to 95%) of pancreatic cancer patients harbor a mutation in K-Ras ^13, 24, 38^. Unfortunately, thus far, no effective inhibitor of the K-Ras oncoprotein has reached the clinic. In this study, we have strived to test the efficacy of DNA2 nuclease inhibition in cancer cells that experience oncogene-induced replication stress. Since such stress is a common denominator among multiple cancer types, DNA2 inhibition could be an efficacious strategy to treat a broad spectrum of cancers. Besides targeting cancer cells with an activated oncogene, chemical inhibition of DNA2 may also be useful for sensitizing cells to DNA damage when combined with radiomimetic chemotherapy or radiation therapy. As we expected, recent studies reported that DNA2 inhibition can sensitize breast cancer cells to chemotherapy inducing DNA damage ^36^. We developed a simple high-throughput screening assay for DNA2 inhibitors, which can screen compound collections in very small reaction volume and in general to screen for inhibitors of other nucleases. We identified several compounds with the ability to inhibit DNA2 in DNA incision and also in a reconstituted system of DNA end resection. The NSC-105808 compound inhibits DNA2 but not other tested nucleases, however additional cellular targets cannot be excluded. Importantly, NSC-105808 treatment recapitulates several phenotypes of cells with DNA2 depletion, such as attenuated DSB end resection, decreased homologous recombination, and suppression of the sensitivity of FANCD2 deficient cells to cisplatin ^31^. DNA2 plays an important role in telomere replication ^35^, thus the effect of NSC-105808 on telomere stability cannot be excluded. The inhibitor reduces the growth of various cancer cells, and specifically those cells in which replication stress is induced via oncogene activation. A similar range of IC_50_ values for the NSC-105808 *in vivo* and in *in vitro* assay may indicate that even minimal inhibition of DNA2 in cells could be lethal to cells. In example a single unrepaired DSB is lethal for cells. Alternatively DNA2 sensitivity to the drug can be altered by cell specific features such as DNA2 posttranslational modifications. Together our work suggests that temporary inhibition of DNA2 nuclease represents a promising strategy in cancer therapy.

## Materials and Methods

### Human cell cultures

U2OS and MCF10A cells were purchased from ATCC and were maintained in McCoy’s 5A medium supplemented with 10% fetal bovine serum (FBS). MCF10A cells were cultured in mammary epithelial growth medium containing insulin, hydrocortisone, epidermal growth factor, and bovine pituitary extract purchased from Clonetics. MCF10A cells with stable H-Ras expression were generated by infection with lentiviral inducible-V5 construct (Life Technologies) and maintained in the presence of G418 (200 µg/ml) and Blasticidin (4 µg/ml). AsPC-1 and PANC-1 cells were cultured in Dulbecco modified Eagle medium (DMEM) (Corning cellgro) with 10% FBS, Vitamin (100×), non-essential amino acids (100×) and PS (100×). BT549 cells were cultured in RPMI-1640 medium with 0.023 IU/ml insulin and Hs578T cells in DMEM medium with 0.01 mg/ml bovine insulin. The pancreatic cell lines (DT-PD59, DT-PD77-E6/E7) were kindly provided by Dr. Michel J. Quellette (University of Nebraska Medical Center) and cultured as described ^10^. HPDE cells with inducible K-Ras expression were generated and maintained in the KSFM medium (Life Technologies) as previously described ^47^. A series of breast epithelial cell lines transformed by H-Ras (MCF10A, MCF10AT1, MCF10DCIS and MCF10CA1a) were obtained from Dr. Isabelle Bedrosian (Department of Surgical Oncology, MD Anderson Cancer Center) and maintained as described ^15^. In the experiments described in Figures 5b and c, MCF10A cells were obtained from ATCC. FANCD2-/- and control cell lines were obtained from Dr. Randy Legerski (Department of Genetics, MD Anderson Cancer Center). Cells were maintained in DMEM containing 10% heat-inactivated FBS and 1% penicillin/streptomycin at 37°C in 5% CO_2_. All cell lines were validated by STR DNA fingerprinting using the AmpF STR Identifiler kit according to manufacturer instructions (Applied Biosystems cat. # 4322288). The STR profiles were compared to known ATCC fingerprints, to the Cell Line Integrated Molecular Authentication database (CLIMA) and to the MD Anderson fingerprint database. The STR profiles matched known DNA fingerprints or were unique.

### Plasmids, interfering RNAs (shRNAs), and transfection

The pLenti6.3/TO/V5-H-Ras vector was obtained from Dr. Shiaw-Yih Lin (Department of Systems Biology, MD Anderson Cancer Center). Briefly, PCMV-H-RAS (V12) was purchased from Invitrogen and subcloned into the pLenti6.3/TO/V5-DEST vector (Invitrogen) based on the commercial protocol of ViraPower TM HiPerform TM T-Rex TM Gateway Expression system. pLenti3.3/TR and pLenti6.3/TO/V5-H-Ras were stably integrated into MCF10A cells. Expression of H-Ras was induced by the addition of Doxycycline (1 μg/ml) for 48 hours. The vector that expresses FLAG-tagged DNA2 has been described ^42^. DNA2 shRNA vectors and control shRNA vectors against luciferase sequences were obtained from Sigma (shDNA2#1: CCGGA CCTGG TGTTG GCAGT CAATA CTCGA GTATT GACTG CCAAC ACCAG GTTTTTTTG; shDNA2#2: CCGGA GTTTG TGATG GGCAA TATTT CTCGA GAAAT ATTGC CCATC ACAAA CTTTTTTTG). On-target smart pool siRNAs against DNA2 were purchased from Dharmacon Research (#9: AGACAAGGUUCCAGCGCCA; #10: UAACAUUGAAGUCGUGAAA; #11: AAGCACAGGUGUACCGAAA; #12: GAGUCACAAUCGAAGGAUA). Transfection of plasmids was performed with the FuGENE 6 reagent from Roche. Transfection of siRNAs was performed with Oligofectamine (Invitrogen).

### Immunohistochemical staining

Tissue microarrays of formalin-fixed, paraffin-embedded (FFPE) pancreatitis, pancreatic tumors with their corresponding nonneoplastic tissues were obtained from US Biomax. FFPE slides of a conditional LSL-K-Ras^G12D^/transgenic pancreatic cancer mouse model was obtained from Dr. Chinthalapally Rao (Department of Internal Medicine, University of Oklahoma). After slides were deparaffinized with xylene and rehydrated with ethanol, antigen retrieval was carried out by placing the slides in 0.1 M sodium citrate (pH 6.0) in a water bath for 30 min at 95°C. Endogenous peroxidase activity was quenched with 3% hydrogen peroxide in PBS for 10 min. Slides were washed three times with PBS and incubated in blocking buffer containing 10% goat serum for 30 min. The slides were incubated with DNA2 antibody (1:200) overnight at 4°C in a wet box, and immunohistochemical staining was done by using the Vectastain ABC avidin biotin-peroxidase enzyme complex kit (Vector Laboratories). Slides were counterstained with hematoxylin and mounted with Permount aqueous medium.

### Cell proliferation assay

Cell proliferation was measured by MTT (3-(4,5-dimethylthiazol-2-yl)-2,5-diphenyltetrazolium bromide; Sigma) reduction (Supplementary Materials and Methods).

### Colony-forming assay

Transfection and colony-forming assay were performed as previously described (^42^, Supplementary Materials and Methods).

### Senescence assay

The assay was performed with a senescence β-galactosidase staining kit (#9860; Cell Signaling) according to the manufacturer’s protocol (Supplementary Materials and Methods).

### Flow cytometric analysis of apoptosis and BrdU incorporation

Apoptosis was measured by flow cytometry using the Annexin V FITC assay kit based on the protocol from the manufacturer (Invitrogen). For BrdU analysis, after treatments, cells were labeled with BrdU (Sigma) for 30 min prior to fixation with 70% cold ethanol (-20°C). The next day, cells were washed with cold PBS and incubated with 2 M HCl for 5 min at room temperature. After adjusting the pH with 0.1 M sodium borate (pH 8.5), cells were incubated in 1% bovine serum albumin with 0.1% Triton X-100 for 30 min and mouse anti-BrdU antibody (1:400) for 1 hr to detect BrdU labeling. After washing, cells were incubated with secondary antibody (fluorescein isothiocyanate 1:400) for 30 min. Cells were resuspended in staining solution (10 μg/mL propidium iodide, 20 μg/mL RNase A, and 0.05% Triton X-100) and cell cycle analysis was performed at the MD Anderson Cancer Center Flow Cytometry and Cellular Imaging Facility. For fluorescence-activated cell sorting (FACS) analysis, the percentage of cells with positive BrdU staining in each phase of the cell cycle was quantitated with the CellQuest software and ModFit software (Verity Software House Inc.)

### Gene conversion and SSA Repair Assays

The DR-GFP, pCAGGS, and pCBASce plasmids were kindly provided by Dr. Maria Jasin (Memorial Sloan-Kettering Cancer Center, New York, NY). U2OS cells containing a single copy of the HR repair reporter substrate DR-GFP were generated as previously described ^41^. SSA reporter cells were kindly provided by Dr. Jeremy Stark (Beckman Research Institute of the City of Hope) ^5^. Details in Supplementary Materials and Methods.

### Immunofluorescent staining and chromatin fractionation

For the detection of DNA-damage-induced p-RPA34 (S4/S8), immunofluorescent staining, and the preparation of chromatin fractions and Western blot analyses were performed as described previously ^41, 42^.

### High throughput screen for inhibitors of DNA2

The screen entailed the use of oligonucleotide (dT)_30_ having a reporter fluorescent moiety, 6-FAM^TM^ (excitation max. = 494 nm, emission max. = 520 nm) on the 5’ end, and the dark quenching group, Iowa Black^®^ FQ (absorption max. = 531 nm), on the 3’ end. Additionally a second dark quencher ZEN^TM^ (N,N-diethyl-4-(4-nitronaphthalen-1-ylazo)-phenylamine, absorption max. = 532 nm) is placed between 9th and 10th base (Figure 2a). Initial screen, it was performed with yeast Dna2 (yDna2) ^40^. Then, the top candidate compounds were tested on hDNA2. Identical reaction conditions were used for assaying the nuclease activity of yDna2 and hDNA2. We note that yDna2 is more active on the ssDNA substrate than hDNA2, thus the experiment with ssDNA substrates were done with 10 nM of hDNA2. The reaction buffer was 20 mM HEPES/NaOH, pH 7.5, 60 mM KCl, 1 mM DTT, 100 µg/ml BSA, and 0.05% Triton X-100. The nuclease assay was initiated by adding MgCl_2_ (3 mM final concentration). T5 nuclease was assayed in the buffer (50 mM potassium acetate, pH 7.9, 20 mM Tris-acetate, 10 mM magnesium acetate and 1 mM DTT) provided by the vendor (NEB). The concentration of the fluorogenic substrate was 0.1 µM. The optimal amount of yDna2, hDNA2 and T5 nuclease was determined by utilizing the Kaleidagraph software to fit the data according to Michaelis Menton Kinetics {V=V_0_+V_max_*x/(Km+x)}. For curve fitting, the maximum increase in fluorescence (upon adding 5 nM of yDna2 or 166 nM of hDNA2 or 10 Units of T5 nuclease) corresponds to V_max_, the background fluorescence (without enzyme) corresponds to V_0_ and “x” is the enzyme concentration (nM). All the compounds were dissolved in DMSO and DMSO alone was used as control; the final concentration of DMSO was at 3.33% for protein-based assay and 0.1% for cell based assay. The Z’ value was calculated as previously described ^49^. Cell proliferation was determined using DAPI staining followed by high-throughput microscopy in a 384 well plate setup. In brief, cells were fixed with formaldehyde, washed, and stained with DAPI and the number of nuclei in each well counted by automated microscopy. The data were analyzed as described ^27^.

### Nuclease substrates

The oligonucleotide PSOL6929 (TTCATGGCTTAGAGCTTAATTGCTGAATCT) was 5’ end-labeled with γ^32^P-ATP (Perkin-Elmer) and T4 polynucleotide kinase (NEB). The 2 kb dsDNA substrate is a fragment of the human DNA2 (hDNA2) gene created by PCR with oligonucleotides PSOL4642 (GATCCTCTAGTACTTCTC) and PSOL6134 (TCTACCTCAAGACTGGTCAG) using the DNA2 expression vector pJD72 ^14^ as template; 50 μCi of α^32^P-dCTP (Perkin-Elmer) was included in the reaction to randomly radiolabel the PCR fragment.

### Nuclease assays

Nuclease assays were performed in buffer R (20 mM Na-HEPES pH 7.5, 1 mM ATP, 0.1 mM DTT, 100 µg/ml BSA, 0.05% Triton-X 100, 2 mM MgCl_2_, 100 mM KCl) and contained 0.5 nM ends (2 kb substrate) or 5 nM ends (the ssDNA oligonucleotide PSOL6929) of DNA. Reactions containing BLM also included an ATP regenerating system consisting of 10 mM creatine phosphate and 50 μg/ml creatine kinase. In control reactions, DMSO was used in place of the inhibitor. Reactions with DNA2 alone were incubated at 30°C, and those with DNA2-BLM-RPA and EXO1 were incubated at 37°C for the indicated times. After the addition of SDS to 0.02%, proteinase K to 0.25 μg/μl, and 0.08% Orange G dye with a final glycerol concentration of 10%, the reaction mixtures were incubated for 5 min at 37°C. Electrophoresis was done in native 10% polyacrylamide gels in TBE buffer (ssDNA substrate) or in 1% agarose gels in TAE buffer (40 mM Tris-acetate, pH 7.4, 0.5 mM EDTA) (2kb substrate). Gels were dried onto DEAE paper on top of Whatman filter paper (GE) and then analyzed in a BioRad Personal Molecular Imager FX phosphorimager. Band intensity was quantified using Image Lab software (BioRad). The fraction of DNA substrate digested was determine by measuring the amount of substrate remaining and normalizing to the total substrate in the negative control lane containing no protein.

## Acknowledgements

The research presented here was supported by CPRIT grant RP140456 to G.I. and G.P., R01 GM080600 to G.I., The University of Texas MD Anderson Cancer Center Duncan Family Institute for Cancer Prevention and Risk Assessment grant to G.P., CPRIT grant RP110532 to C.C.S, NIH grants ES015632 and ES015252 to P.S. The authors thank Dr. Kenneth L. Scott (Baylor College of Medicine) for providing K-Ras-HPDE cells.

Supplementary Information accompanies the paper on the Oncogenesis website (http://www.nature.com/oncsis)

